# Effects of parental verbal abuse experience on the Glutamate response to swear words in the ventromedial prefrontal cortex: A functional ^1^H-magnetic resonance spectroscopy study

**DOI:** 10.1101/2022.10.10.511658

**Authors:** Jae Hyun Yoo, Young Woo Park, Dohyun Kim, HyunWook Park, Bumseok Jeong

## Abstract

Several lines of evidence indicate verbal abuse (VA) critically impacts the developing brain; however, whether VA results in changes in brain neurochemistry has not been established in humans. Here, we hypothesized that exposure to recurrent parental VA elicits heightened glutamate (Glu) responses during the presentation of swear words, which can be measured with functional magnetic resonance spectroscopy (fMRS). During an emotional Stroop task consisting of blocks of color and swear words, metabolite concentration changes were measured in the ventromedial prefrontal cortex (vmPFC) and the left amygdalohippocampal region (AMHC) of healthy adults (14 F/27 M, 23±4 years old) using fMRS. The dynamic changes in Glu and their associations with the emotional state of the participants were finally evaluated based on 36 datasets from the vmPFC and 30 from the AMHC.

A repeated-measures analysis of covariance revealed a modest effect of parental VA severity on Glu changes in the vmPFC. Furthermore, the total score on the Verbal Abuse Questionnaire by parents (pVAQ) was associated with the Glu response to swear words (Δ*Glu*_*Sw e*_). The interaction term of Δ*Glu*_*Sw e*_ and baseline N-acetyl aspartate (NAA) level in the vmPFC could be used to predict state-trait anxiety level and depressive mood. We could not find any significant associations between Δ*Glu*_*Sw e*_ in the AMHC and either pVAQ or emotional states.

We conclude that parental VA exposure in individuals is associated with a greater Glu response towards VA-related stimuli in the vmPFC and that the accompanying low NAA level may be associated with anxiety level or depressive mood.

## Introduction

Emotional abuse during childhood can cause significant harm to the child’s development and exert a deleterious effects on adult life (Hart et al., 1997). Among various types of emotional abuse, verbal abuse (VA) is highly prevalent during childhood and adolescence (Hawker and Boulton, 2000). Victims of VA during childhood have been associated with increased psychiatric symptoms such as depression, anxiety, suicidal ideation and even psychosis (Hawker and Boulton, 2000; Schreier et al., 2009; Miller et al., 2017).

Previous literature has suggested a detrimental effect of VA on widespread regions of the developing brain. In a systematic review, childhood maltreatment in subjects was associated with structural alterations in the ‘threat-detection and response circuit’ including the anterior cingulate cortex (ACC), ventromedial prefrontal cortex (vmPFC), hippocampus, thalamus and sensory cortices (Teicher et al., 2016). In parallel with these findings, decreased N-acetyl aspartate (NAA), which is a brain metabolite reflecting neuronal density, has been found in subjects with posttraumatic stress disorder (PTSD) (Ham et al., 2007) and a history of maltreatment (De Bellis et al., 2000). A putative mechanism for this abnormal brain development is an increased response of the hypothalamic-pituitary-adrenal axis to emotional stress (Sale, 2016). Animal models of chronic stress have also revealed that Glu release can facilitate neuronal remodeling in the PFC-limbic circuit and working memory impairment in rodents (Nathan et al., 2004; Mitra et al., 2005).

The vmPFC serves as a hub in the default mode network and has broad functional roles in affect regulation, self-reference, pain and visceral sensation (Roy et al., 2012). It has been shown to have a high metabolism rate in a resting state (Raichle et al., 2001) but attenuated activity during goal-directed behavior (Harrison et al., 2011). This task-induced deactivation in the vmPFC has been observed in functional neuroimaging studies such as those using an emotional face-matching task (Rest – Shape or Faces) or the color-word Stroop task (Rest – Congruent or Incongruent condition). In a Macaque monkey study using positron emission tomography, a higher level of regional blood flow in the vmPFC was found during spatial working memory task, and such task-induced deactivation was associated with better performance (Kojima et al., 2009).

In contrast, activation of the vmPFC has been consistently reported in functional magnetic resonance imaging (fMRI) studies when subjects were exposed to a negative emotional stimulus (Wager et al., 2008; Lindquist et al., 2012). According to a neurocircuitry model of emotion processing, recruitment of the vmPFC downregulates amygdala activity to modulate the negative affective response (Etkin et al., 2011). Failures in negative emotion regulation in PTSD subjects have been associated with a hyperresponsive amygdala and hyporesponsive vmPFC in response to traumatic scripts (Hughes and Shin, 2011). An fMRI study using the Stroop task also showed reduced deactivation in women with childhood abuse during the processing of threatening words compared to that during the processing of positive words (Mackiewicz Seghete et al., 2017). These findings suggest that the vmPFC plays a pivotal role as a mediator of emotion regulation. However, significant gaps exist between maltreatment-related Glu changes within the emotion processing circuit and emotional states due to the lack of functional magnetic resonance spectroscopy (fMRS) studies.

We hypothesized that exposure to VA-related stimuli would elicit negative emotion and that regulation of the emotional response might result in an increase or maintenance of the Glu concentration in the vmPFC and nonsignificant changes in Glu in the amygdalohippocampal region (AMHC). Alternatively, a simple cognitive task might induce deactivation of the vmPFC, indicated by a decrease in the Glu concentration. Dynamic Glu changes during emotion regulation might be affected by histories of parental VA. Finally, neurochemical profiles and parental VA severity might be associated with emotional states such as anxiety or depressive mood. To explore those relationships, we acquired fMRS signals during performance on an emotional Stroop task consisting of three types of blocks: at rest, a color-word Stroop task, and emotional interference Stroop task.

## Methods

### Participants and screening

Forty-three healthy young adults were recruited (15 females; 23.34 ± 3.93 years of age, range 18-35 years of age). Detailed information regarding the study was provided to all participants, and written informed consent was obtained prior to enrollment. This study was approved by the Institutional Review Board at Korea Advanced Institute of Science and Technology (KAIST: KH2017-058). Following consent, all participants were interviewed using the Korean version of the Diagnostic Interview for Genetic Studies (DIGS) (Joo et al., 2004) by two psychiatrists (D Kim, JH Yoo) for screening of psychiatric disorders. After the screening interview, two participants were excluded due to a lifetime history of depression.

We measured the 15 types of VA experience using the Korean version of the Verbal Abuse Questionnaire (pVAQ) (Jeong et al., 2015) including scolding, yelling, swearing, blaming and threatening from parents (Teicher et al., 2010; Jeong et al., 2015). The participants were also given self-report questionnaires for the Center for Epidemiological Studies Depression Scale (CES-D) (Radloff, 1977; Cho and Kim, 1998) and State-Trait Anxiety Inventory (STAI) (Hahn, 1996; Spielberger et al., 2017), all translated to Korean, to screen their emotional state.

The exclusion criteria of participants were (1) an intelligence quotient (IQ) < 70, (2) a current/past history of brain trauma, organic brain disorders, seizure or any neurological disorders, (3) any psychiatric disorder history according to the Diagnostic and Statistical Manual of Mental disorder, 5th Edition (DSM-5), and (4) a history of traumatic experience other than verbal and emotional abuse.

### Experimental design

The current study used a modified version of the emotional Stroop task (Lee et al., 2017). Our paradigm consisted of five alternating blocks: an initial rest period (rest 1, 4 minutes), a facilitation Stroop task with words representing color for minimal emotional arousal (color block, 3 minutes), a second rest period (rest 2, 5 minutes), a Stroop task with swear words for negative emotional arousal (swear block, 3 minutes) and a final rest period (rest 3, 5 minutes). In the facilitation Stroop block, three congruent color words (Blue, Red, and Yellow) were repeatedly presented to participants to explore Glu changes elicited by simple cognitive process without any interference of emotion. In contrast, we chose 20 swear words for the emotional Stroop block and colored them with blue, red, and yellow to evoke a strong emotional interference-associated brain response similar to that associated with VA exposure. Swear words (emotional valence, −3.078 ± 0.636) were selected from the 240 commonly used Korean words whose valence (Likert scale from −5 to 5) was rated from a satellite group of 30 participants (10 females; age range 23-33 years) prior to the fMRS experiment. We placed the facilitation block ahead of the emotional block to ensure the participant’s mood was not disturbed by the swear word stimulus presentation. However, this block order could create a conditioning effect for both the color and emotional Stroop task during the second voxel of interest (VOI) acquisitions. To mitigate this effect, the word stimulus was presented to the participant using a different pseudorandom order for each VOI. In addition, the fMRS spectra from each VOI were acquired in a different order among participants to further minimize the conditioning effect in group statistics.

Each Stroop block contained 60 different word stimuli that were presented for one second, with a two-second-long fixation cross presented between the stimuli. Five minutes of rest blocks were placed following each 3-minute task block to ensure the Glu levels were back to stable conditions following the functional activations. The acquisition of fMRS data took place for 20 minutes per VOI (Figure 1). Participants were asked to match the exact color of the words using a 3-button keypad.

**Figure 1.**
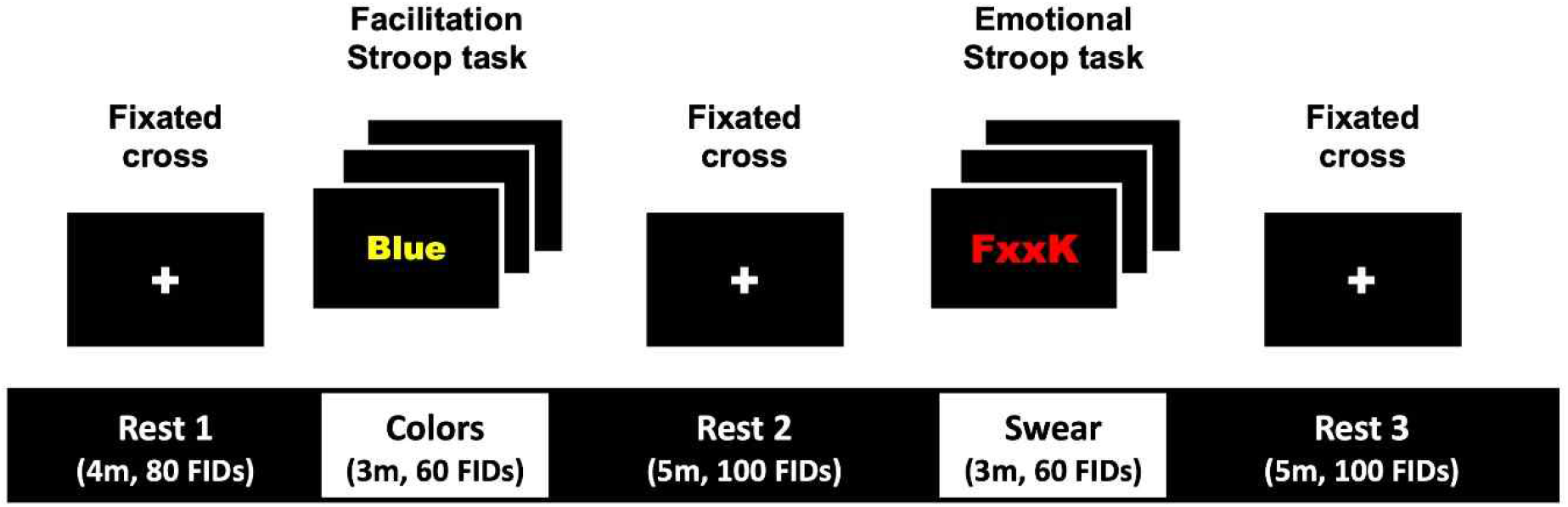

### Data acquisition and postprocessing of fMRS data

*In vivo* data acquisitions were performed using the Siemens 3T Verio scanner (Erlangen, Germany) at KAIST using a 32-channel radio frequency (RF) coil. T1-weighted magnetization-prepared rapid gradient echo (MPRAGE) sequences (repetition time (TR)/ inversion time (TI)/ echo time (TE) = 2400/1000/2.02 ms; 0.7-mm^3^ resolution) were acquired at the beginning of the study for both fMRS VOI prescription and anatomical analysis. We placed the vmPFC voxel primarily in Brodmann area (BA) 32 with a volume of 6 cc (LRxAPxFH = 20×15×20 mm^3^). The AMHC voxel had a slightly smaller volume at ~4 cc (LRxAPxFH = 15×22×12 mm^3^) and was placed over regions of BA 34 and 28 (Figure 2). Each VOI was prescribed after review of its location by two or more trained experts. The Montreal Neurological Institute (MNI) coordinates of the center of gravity of each VOI were x=0.7, y= 44.2, z=-3.5 for the vmPFC and x=-28.3, y=-15.4, z=-18.1 for the left AMHC.

**Figure 2.**
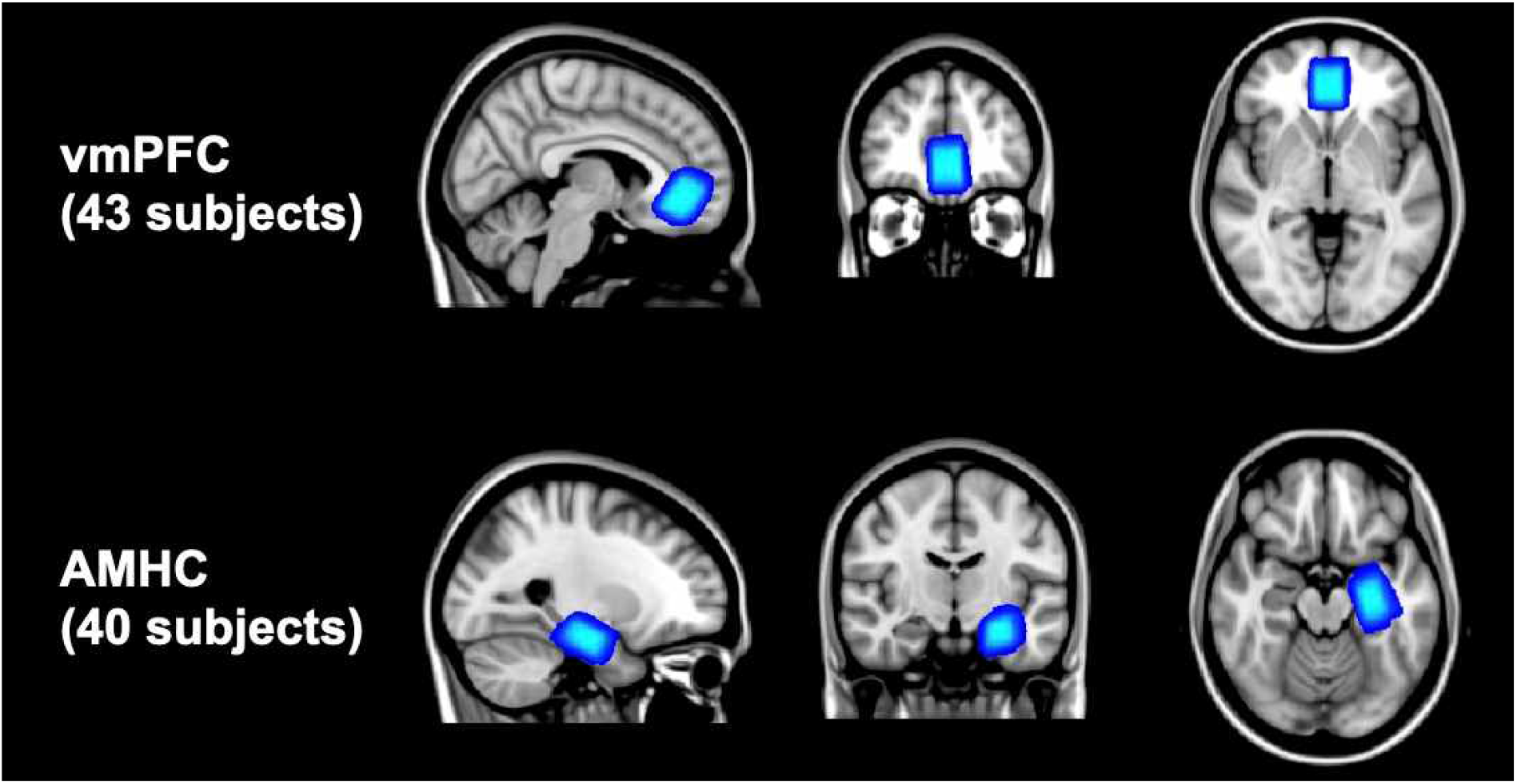

B_0_ shimming was performed using FAST(EST)MAP (FM) (Gruetter and Tkac, 2000), and MR spectra were acquired using the modified semi-LASER (sLASER) sequence (Oz and Tkac, 2011) (TR/TE = 3000/28 ms). The semi-LASER sequence has advantages as it maintains a uniform B1 field and a desired flip angle within voxels relative to the PRESS sequence (Zhu and Barker, 2011). Following FM, the RF power for the 90º excitation and the VAPOR water suppression (WS) pulses were calibrated. Spectra were acquired with a 70-Hz WS bandwidth (BW), a sampling BW of 6 kHz and 2048 complex points. Additional unsuppressed water scans were obtained for eddy current correction and metabolite quantification, as described previously (Deelchand et al., 2015). Four hundred single-shot spectra were acquired during a 20-minute acquisition period, and the eddy current, frequency and phase variations of the acquired spectra were corrected by using MRspa (Deelchand, 2016).

### Metabolite estimation using the linear combination (LC) model and data quality analysis

Prior to the acquisition, a single-shot spectrum of the unsuppressed water signal was measured to measure field homogeneity, and datasets with water peak linewidths of 10 Hz or greater were excluded from the analysis. In addition, each individual spectra for each participant was screened for any underlying coherence artifacts. Based on the spectral quality inspection, two vmPFC and eight AMHC datasets were found to be of insufficient quality and were excluded from the statistical analysis. Finally, we excluded datasets from three participants who had slept or failed to respond to five or more stimuli during the functional data acquisition. The data quality screenings resulted in 36 vmPFC and 30 AMHC remaining datasets, which were used for the statistical analysis.

Averages of 60 free induction decays (FIDs) from each of the five event blocks were summed for each participant. For event blocks with more than 60 FIDs, such as rest 2 and 3, we summed 60 FIDs from the end to minimize the effect of the preceding event block. Metabolite quantification of the summed sLASER spectra were performed with LCModel 6.3-0G (Provencher, 1993) with the water-scaling option (Deelchand et al., 2015). The fraction of cerebrospinal fluid (CSF) within the VOI was estimated by first generating 3D masks for each VOI prescription from the file header of the corresponding fMRS data and then applying the masks on the 3D tissue map generated by the tissue segmentation of T1-weighted data with SPM12 (Ashburner and Friston, 2007). Metabolites with Cramer-Rao lower bounds (CRLB) greater than 20% were excluded in the neurochemical analysis.

### Statistical analysis of metabolite estimations

Evidence from a previous fMRI study suggested that functional changes in frontolimbic networks during negative emotion processing were significantly associated with depressive symptoms as well as previous VA experiences (Lee et al., 2015). Likewise, Glu concentrations in the vmPFC or AMHC may change according to brain activation, and two major contrasts are cognitive effort during a task and implicit emotion processing. To verify our hypothesis, we first calculated percent changes in Glu concentrations during swear (Δ*Glu*_*Sw e*_) and color (Δ*Glu*_*Cobr*_) blocks compared to each preceding rest block and then analyzed those variables using one-way repeated measures analyses of variance (RM-ANOVA). In addition, we conducted an exploratory repeated measures analyses of covariance (RM-ANCOVA) to evaluate a possible moderating effect of pVAQ on the changes in Glu concentrations. Post hoc paired comparisons were also performed between two different stimulus blocks and between each stimulus block and the preceding rest block to detect any meaningful differences between the event blocks.

Second, the impact of parental VA exposure on those brain metabolite levels that reflect functional and structural changes was examined. Relationships between pVAQ and neurochemical markers such as Δ*Glu*_*Sw e*_ and baseline NAA (bNAA) were analyzed using Pearson’s Correlation analysis. We further examined whether pVAQ score was associated with either Δ*Glu*_*Sw e*_ or Δ*Glu*_*Cobr*_.

Next, we speculated that the pVAQ and Glu responses could explain subjective emotional state such as anxiety and depression. To find the best predictors of emotional state (STAI-state anxiety (S), STAI-trait anxiety (T), and CES-D), we first proposed the stress-susceptibility model [Model 1] in which pVAQ might have a direct association with emotional states. Alternatively, we proposed that a metabolite model ofΔ*Glu*_*Sw e*_ against bNAA, representing a Glu change per neuronal density, could explain subjective emotion [Model 2]. We explored significant predictors by following the three linear models and comparing the performance of each model.

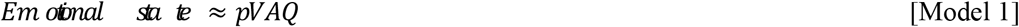

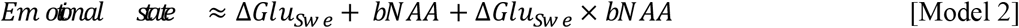

We also compared the model fitness among the proposed models to find the best model and contributing factors and conducted RM-ANOVA analysis for other key identifiable metabolites such as NAA, creatine (Cr), choline (Cho) and myo-inositol (Ins) to examine dynamic changes during the emotional Stroop task.

Statistical analyses were performed using R software version 3.5.0 (R Core Team, 2013). Factors in the linear models were normalized within participants, and their main effects were controlled for age and gender as covariates.

## Results

### Sample characteristics

The demographic and clinical characteristics of participants are described in Table 1. The ethnic background of all participants was Korean. Based on the criteria used in a previous study (Jeong et al., 2015), 1 participant had been exposed to moderate parental VA (pVAQ range of 20-42), 19 to mild (7-19), and 16 to minimal (0-6). Participants had relatively low STAI-S and STAI-T scores ranging from 20 to 57 and 20 to 56, respectively. Total CES-D scores ranged from 0 to 23 among the group, and only three participants had probable depressive symptoms (CES-D range of 16-25).

**Table 1.** Participant Characteristics

|  | vmPFC (n=36) | AMHC (n=30) |
| --- | --- | --- |
| Age, y | 23.06 ± 3.54 | 22.99 ± 3.34 |
| Gender (Male : Female) | 23:13 | 17:13 |
| Full scale IQ | 123.67 ± 11.11 | 122.17 ± 11.31 |
| pVAQ | 7.43 ± 6.21 | 7.47 ± 6.51 |
| CES-D | 8.14 ± 5.60 | 7.73 ± 5.35 |
| STAI-S | 35.25 ± 8.22 | 34.77 ± 8.55 |
| STAI-T | 35.22 ± 8.17 | 34.73 ± 8.37 |
| Tissue proportion (%) |  |  |
| GM | 75.69 ± 3.09 | 71.62 ± 7.14 |
| WM | 13.83 ± 3.76 | 23.22 ± 7.17 |
| CSF | 10.47 ± 2.24 | 5.16 ± 1.95 |
| SNR estimated by LCModel | 31.03 ± 7.52 | 12.97 ± 2.68 |
| Linewidth at baseline | 7.19 ± 1.08 | 7.94 ± 0.77 |
| Linewidth after task | 7.89 ± 1.37 | 8.23 ± 0.95 |

pVAQ, Verbal Abuse Questionnaire – parental origin; CES-D, Center for Epidemiological Studies Depression Scale; STAI-S, State-Trait Anxiety Inventory – State anxiety; STAI-T State-Trait Anxiety Inventory – Trait anxiety; GM, gray matter; WM, white matter; CSF, cerebrospinal fluid; SNR, signal-to-noise ratio

### Metabolite quantification results

The LC Model fitting results across different metabolites are depicted in Figure 3. Five metabolites (Glu, NAA, Cho, Cr, Ins) and macromolecules were reliably quantified (CRLB < 20%) in both the vmPFC and AMHC.

**Figure 3.**
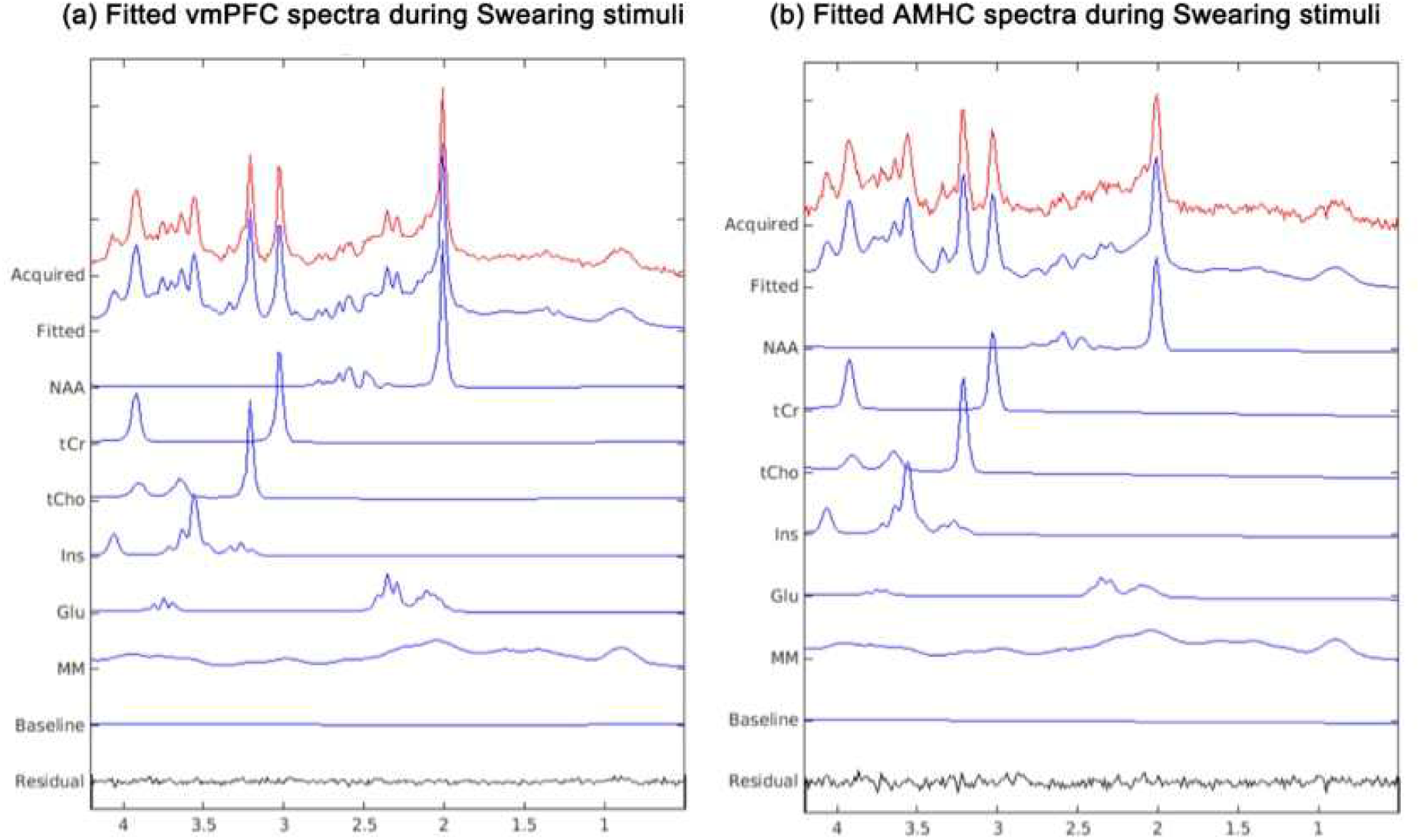

### Task and emotion effects on Glu levels in the vmPFC and AMHC

An exploratory one-way RM-ANOVA to observe effects of emotion on Glu concentration in the vmPFC did not yield significant results (F(1,35) = 2.97, p = 0.094). However, after controlling individual parental VA exposure severity with an RM-ANCOVA model, the effect of emotion on Glu concentration was marginally significant (F(1,34) = 4.12, p = 0.050). A post hoc paired comparison demonstrated a significant decrease in Glu concentration during the color block compared to that during the preceding rest block (t(35) = 2.43, p = 0.020) and swear block (t(35) = 3.02, p = 0.005). However, the Glu changes in the swear block did not differ from those in the preceding rest block (t(35) < 0.01, p = 0.994, Figure 4).

**Figure 4.**
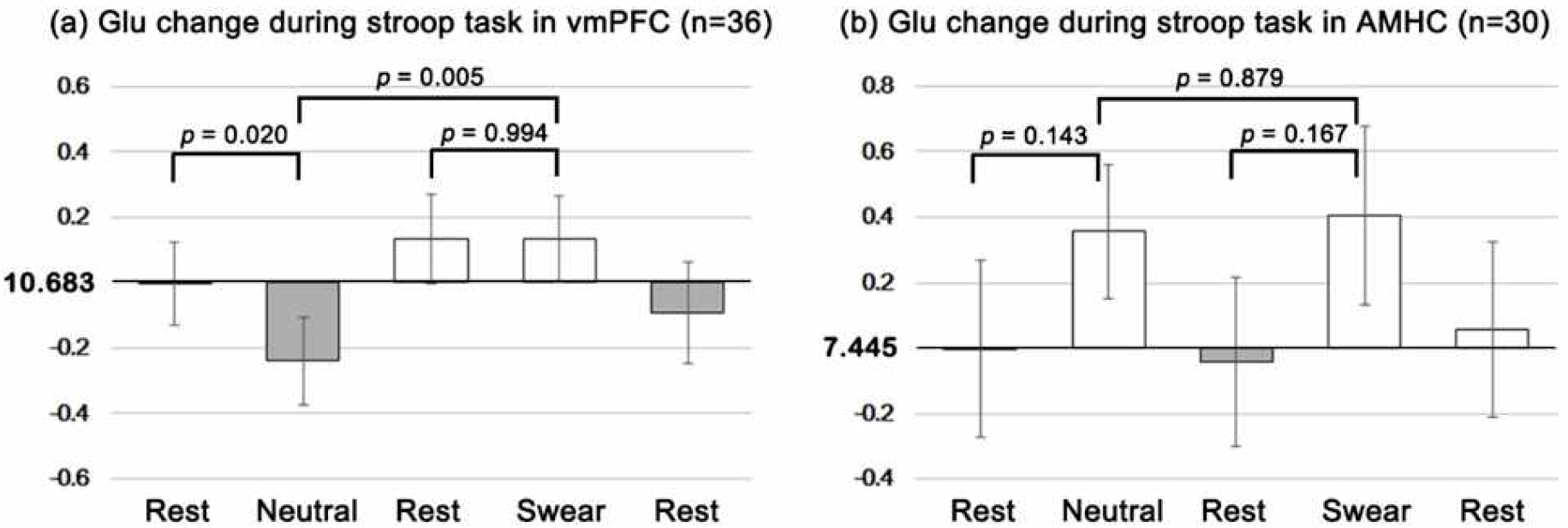

One-way RM-ANOVA of AMHC Glu revealed that Δ*Glu*_*Sw e*_was not significantly higher than Δ*Glu*_*Cobr*_ (F(1,35) = 0.02, p = 0.900). An emotion effect was still absent even after controlling for the pVAQ score (F(1,34) = 0.05, p = 0.826). Post hoc paired comparison analysis showed no significant difference in Glu concentration in the AMHC when comparing the AMHC Glu concentration in the swear block to that in rest 2 (t(29) =-1.42, p=0.167), the AMHC Glu concentration in the color block to that in rest 1 (t(29) =-1.51, p=0.143), or the AMHC Glu concentration in the swear block to that in the color block (t(29) =0.15, p=0.879).

### Association among parental VA exposure, metabolite changes and emotional state

The correlation analysis showed a positive association between pVAQ score and Δ*Glu*_*Sw e*_ (Pearson’s r= 0.36, p = 0.031, Figure 5a) but not between pVAQ score and bNAA (Pearson’s r= −0.12, p = 0.488, Figure 5b) in the vmPFC. In addition, the correlation between Δ*Glu*_*Sw e*_ and bNAA in the vmPFC approached the trend level of significance (Pearson’s r= −0.29, p = 0.086). In the AMHC, however, no significant correlations were found between pVAQ and *Glu*_*Sw e*_ (Pearson’s r= −0.01, p = 0.945), pVAQ and bNAA (Pearson’s r= −0.10, p = 0.598), or Δ*Glu*_*Sw e*_ and bNAA (Pearson’s r= −0.31, p = 0.096). In the linear model analysis, pVAQ was a notable predictor of Δ*Glu*_*Sw e*_, which showed a positive association (estimates = 0.34, t(32) = 2.11, p = 0.043), while no significant association was shown between Δ*Glu*_*Cobr*_ and pVAQ (t(32) = −0.46, p = 0.650) in the vmPFC. Neither Δ*Glu*_*Sw e*_ nor Δ*Glu*_*Cobr*_ showed a significant association with pVAQ in the AMHC (t(26) = −0.12, p =0.904 and t(26) = −0.20, p= 0.846, respectively).

**Figure 5.**
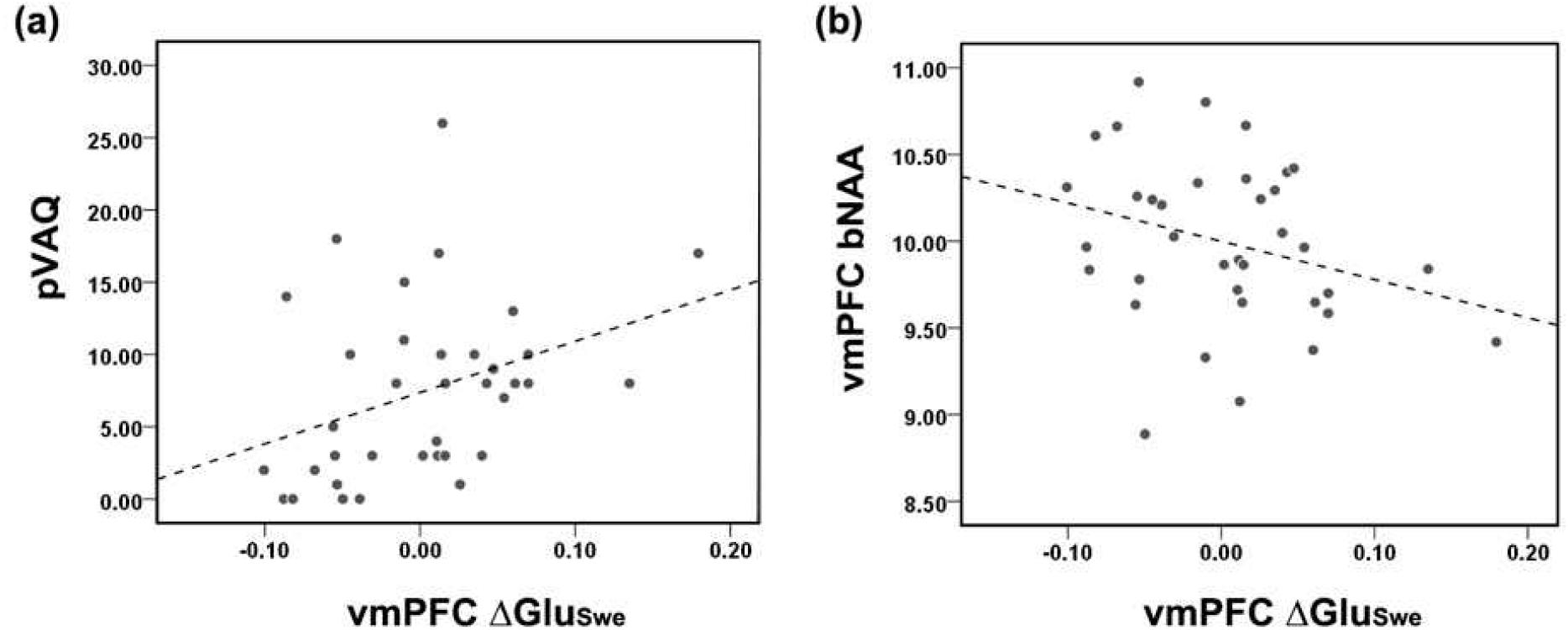

A comparison of the different models showed that the metabolite model (F(5,30) = 3.16, p = 0.057) was marginally superior to the stress-susceptibility model (Table 2) in predicting the STAI-S scores among participants. Significant predictors for STAI-T were pVAQ score in the stress-susceptibility model (t(32) = 2.28, p = 0.029) and vmPFC Δ*Glu*_*Sw e*_ x bNAA (t(30) = −2.54, p=0.017) in the metabolite model. When explaining CES-D scores, the only significant predictor was vmPFC Δ*Glu*_*Sw e*_ x bNAA in the metabolite model (t(30) = −2.67, p = 0.012).

**Table 2.**
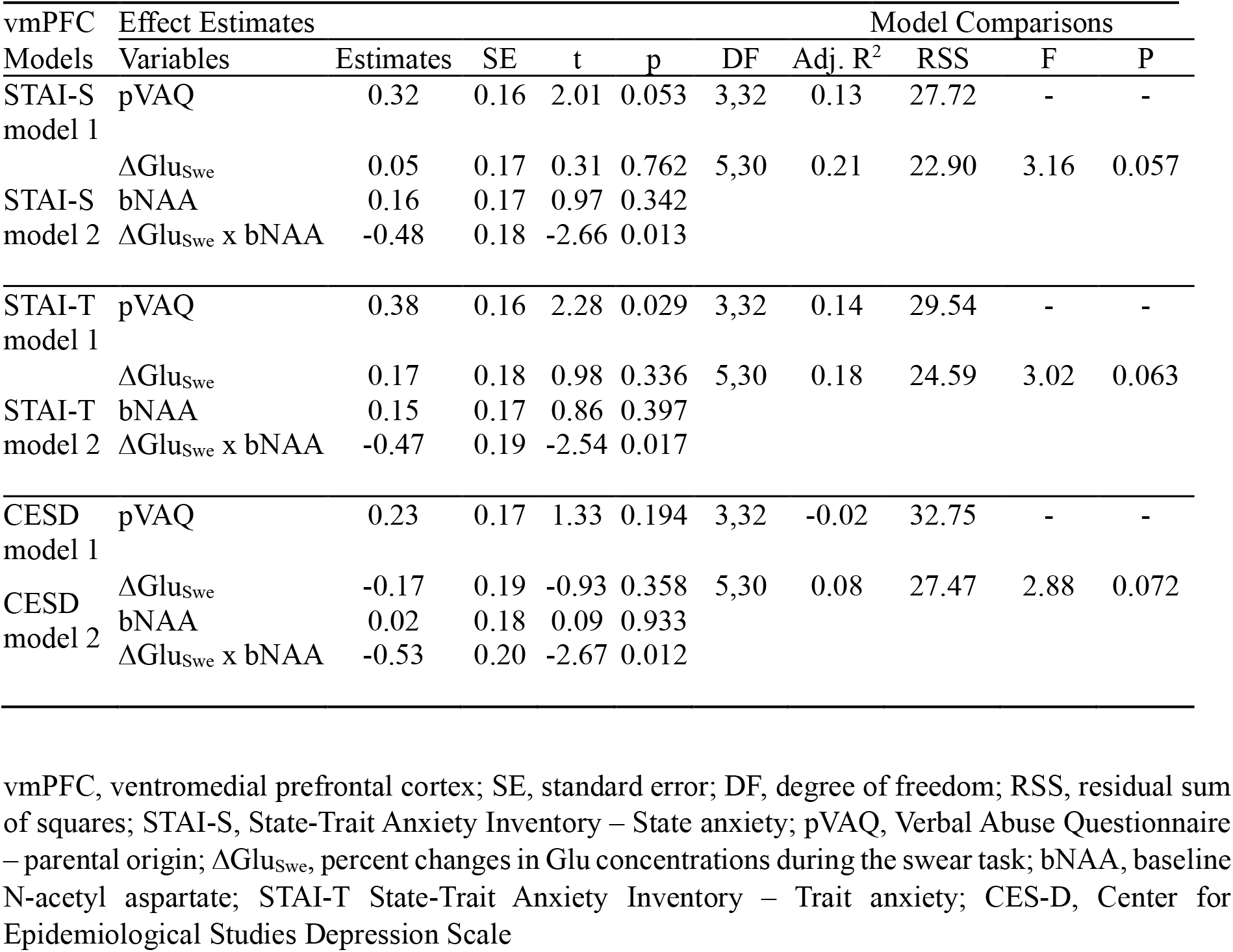
Predictors from stress-susceptibility and vmPFC metabolites explaining emotional states

Among the 30 AMHC datasets, no significant association was found for pVAQ with Δ*Glu*_*Sw e*_, bNAA orΔ*Glu*_*Sw e*_ x bNAA in any proposed models explaining STAI-S, STAI-T and CES-D scores. (Table 3).

**Table 3.**
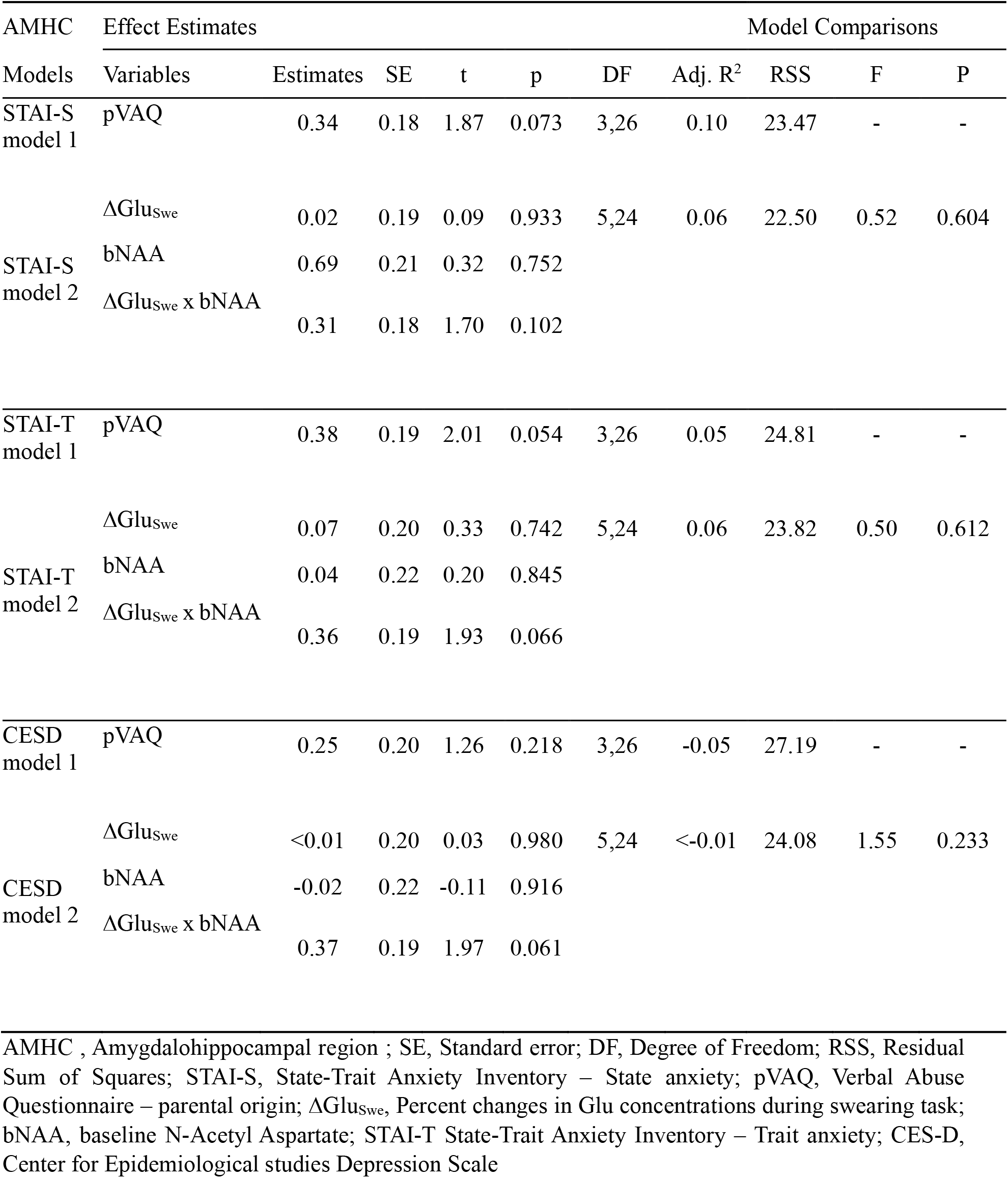
Predictors from stress-susceptibility and AMHC metabolites explaining emotional states

### Analysis of other metabolites

Exploratory RM-ANOVA and RM-ANCOVA were also conducted for the metabolites other than Glu (NAA, Cho, Cr, Ins). However, metabolites from neither the vmPFC nor the AMHC were significantly modulated according to emotional interference or pVAQ severity (Table 4).

**Table 4.** One-way repeated measures ANOVA and ANCOVA results for metabolites other than glutamate

| Metabolites | Results of one-way RM-ANOVA and RM-ANCOVA |  |  |  |  |
| --- | --- | --- | --- | --- | --- |
|  | Model | SS(Type III) | DF | F | p-value |
| vmPFC NAA | RM-ANOVA | < 0.01 | 1,35 | 0.71 | 0.406 |
|  | RM-ANCOVA | < 0.01 | 1,34 | < 0.01 | 0.962 |
| AMHC NAA | RM-ANOVA | < 0.01 | 1,29 | 0.46 | 0.504 |
|  | RM-ANCOVA | < 0.01 | 1,28 | < 0.01 | 0.958 |
| vmPFC Cho | RM-ANOVA | < 0.01 | 1,35 | 0.02 | 0.899 |
|  | RM-ANCOVA | < 0.01 | 1,34 | 0.98 | 0.328 |
| AMHC Cho | RM-ANOVA | < 0.01 | 1,29 | 0.19 | 0.665 |
|  | RM-ANCOVA | 0.01 | 1,28 | 1.20 | 0.282 |
| vmPFC Cr | RM-ANOVA | <0.01 | 1,35 | 0.22 | 0.644 |
|  | RM-ANCOVA | < 0.01 | 1,34 | 0.13 | 0.723 |
| AMHC Cr | RM-ANOVA | 0.02 | 1,29 | 2.04 | 0.164 |
|  | RM-ANCOVA | < 0.01 | 1,28 | 0.13 | 0.720 |
| vmPFC Ins | RM-ANOVA | < 0.01 | 1,35 | 0.01 | 0.925 |
|  | RM-ANCOVA | < 0.01 | 1,34 | 0.43 | 0.515 |
| AMHC Ins | RM-ANOVA | < 0.01 | 1,29 | 0.05 | 0.819 |
|  | RM-ANCOVA | < 0.01 | 1,28 | 0.14 | 0.715 |

RM-ANOVA, repeated measures analysis of variance; RM-ANCOVA, repeated measures analysis of covariance; SS, sum of squares; DF, degree of freedom; NAA, N-acetyl aspartate; Cho, choline; Cr, creatine; Ins, myo-inositol

## Discussion

In the current study, we examined changes in Glu concentrations from the two key regions related to processing emotions, the vmPFC and left AMHC, in healthy young adults with various levels of parental VA exposure. Our RM-ANCOVA findings revealed that the task-induced decrease in Glu concentrations within the vmPFC but not the left AMHC was modulated by an emotional interference effect and pVAQ score. This modulation is supported by findings from correlation analyses showing that pVAQ was significantly associated with Δ*Glu*_*Sw e*_ but not Δ*Glu*_*Cobr*_ in the vmPFC. In addition, vmPFC Δ*Glu*_*Sw e*_ x bNAA was a crucial factor that predicted state and trait anxiety and depressive mood, while the pVAQ score was only significantly associated with STAI-T score. However, neither emotion interference nor pVAQ were found to affect Glu changes in the AMHC, and the linear model results did not reveal any significant association between pVAQ and AMHC Δ*Glu*_*Sw e*_ or their supporting role in predicting individual emotion states.

Previous studies have postulated that the vmPFC and AMHC regions play a key role in the ‘threat-detection and response circuit’ that may be modified in response to abusive experience (Teicher et al., 2016). Several studies examining neurobiological correlates of abuse have suggested that threat-related neural responses are first processed at the AMHC, followed by the vmPFC during the regulation of emotion (Myers-Schulz and Koenigs, 2012; Motzkin et al., 2015). Indeed, atrophic changes in the hippocampus and vmPFC regions have been associated with maltreated individuals (Vythilingam et al., 2002; Kelly et al., 2013; Chaney et al., 2014). The inferred controllability information was leveraged to increase motivated behavior in the vmPFC (Kim et al., 2021). Expecting social punishment such as swear words facilitates control over a decision under uncertainty by recruiting medial prefrontal cortex (Kim and Jeong, 2020).

Herein, we performed fMRS studies of two hub regions of the ‘threat-detection and response circuit’ to gain novel insights into the detrimental effects of parental VA experience. The current experiment revealed a particular association of pVAQ with Glu response to VA-related emotional interference in the vmPFC. Another noteworthy finding is that the changes in metabolite concentrations within the vmPFC may be used to predict individual state-trait anxiety level and measure the variance of depressive mood. Finally, our study demonstrated the feasibility of reliably acquiring fMRS signals from these two VOIs, which is a challenge due to the severe magnetic susceptibility effect near air-tissue boundaries. We were able to consistently obtain high-quality fMRS signals from each VOI over a large number of subjects using advanced spectroscopy sequences and shimming techniques, demonstrating the future feasibility of integrating fMRS scans in standard 3T clinical scanners.

According to close examination of the anatomical and functional connectivity patterns, the ACC region has been hypothetically divided into two parts, the dorsal ‘cognitive’ division and the ventral ‘affective’ division (Bush et al., 2000). The vmPFC VOI in the current study was placed in the affective division across the pregenual ACC (pgACC) and subgenual ACC (sgACC). A growing body of evidence suggests that the pgACC plays a special role in emotional regulation, autonomic integration, and affect related to pain (Myers-Schulz and Koenigs, 2012; Manohar and Husain, 2016; Hiser and Koenigs, 2018). Imaging studies have demonstrated that tasks associated with the appraisal of negative affect or regulation of emotional interference consistently elicited activation of both the pgACC and sgACC (Zald et al., 2002; Egner et al., 2008; Etkin et al., 2011). However, studies using the cognitive Stroop task showed deactivation of vmPFC regions in the presence of engagement of the dorsal ACC (Peterson et al., 1999; Leung et al., 2000). Additionally, the pgACC and its neuroanatomical network is also a key contributor for obtaining and evaluating information about the social context and relate this information to structures involved in emotion generation (Stevens et al., 2011). A meta-analysis of neuroimaging studies related to social exclusion, rejection, and negative evaluation suggested a robust role of the pgACC and sgACC in social pain elicitation and self-reported distress (Rotge et al., 2015). For successful performance in our color-naming task in the emotional Stroop block, the vmPFC might engage to minimize both the emotional interference effect of word stimuli and their previous VA exposure. In this perspective, the heightened Glu response to swear word stimuli in the vmPFC and its association with parental VA exposure might reflect a long-term detrimental effect of the social context created by the VA exposure, although a general influence of the negative emotional context rather than the swear word-specific context cannot be excluded in this study.

Furthermore, we found that the vmPFC Δ*Glu*_*Sw e*_ x bNAA was negatively associated with state-trait anxiety and depressive mood state, while the pVAQ score was only associated with high trait anxiety, suggesting that the effect of vmPFC Δ*Glu*_*Sw e*_ on emotional state was remarkable in individuals with low bNAA. Several ^1^H-MRS studies have suggested that NAA is an indicator of neuronal density, and decreased NAA in pregenual ACC regions has been found in subjects with PTSD (Ham et al., 2007) In addition, structural deficits in vmPFC regions has repeatedly been found in individuals who have experienced maltreatment (Kelly et al., 2013; Chaney et al., 2014). Although we could not find a clear relationship between bNAA and pVAQ in the current study, our results suggest that a heightened neural response and decreased neuronal density of the vmPFC in individuals with parental VA experience could be crucial neurobiological markers predicting anxiety or depressive mood. Using our observations together with previous fMRI evidence (Lee et al., 2015; Lee et al., 2017), we illustrated a conceptual framework of a proposed neurobiological mechanism associated with emotional interference processing in the vmPFC in Figure 6.

Studies examining the neurobiological correlates of abuse have suggested a central role of the amygdala during threat-related neural responses (Myers-Schulz and Koenigs, 2012; Motzkin et al., 2015); however, we were unable to find any significant metabolite changes or factors clearly associated with emotional states in our analysis of the AMHC. A possible explanation for these findings is that our AMHC VOI was small (below 4 cc) and encompassed broad surrounding regions other than gray matter, resulting in a low signal-to-noise ratio (SNR) relative to that in the vmPFC VOI. A separate body of ^1^H-MRS research also reported low SNR in the amygdala region because of the small voxel size and field inhomogeneity due to the proximity of an air-tissue boundary and large vessels (O’Brien et al., 2010; Boubela et al., 2015; Schubert et al., 2017). However, there were some promising findings, such as the marginal association of AMHC Δ*Glu*_*Sw e*_ x bNAA with STAI-S and CES-D, which warrants further fMRS investigation with additional signal averaging, a large sample size including a clinical population, or high-field scanner to reveal neurochemical dynamics relevant to VA experience and its clinical implications in the AMHC.

There are several limitations in this study. First, the total paradigm length was quite long (20 minutes), and several subjects had minor head movements during the scans, particularly during the final 5 minutes of the rest 3 block. We believe that this limitation can be rectified by having an improved task paradigm that evokes and maintains the same level of attention throughout the entire experiment. The use of head motion tracking with a special pulse sequence to compensate for such movements could also alleviate this issue. Second, the comparison of the Glu level between the facilitation color Stroop task and emotional Stroop task might include a cognitive interference (nonmatched color-word) effect as well as emotional interference effect. While prior studies using the cognitive Stroop task showed deactivation of the pregenual ACC, the current results demonstrated differential patterns of the Glu response according to emotional valence (Peterson et al., 1999; Leung et al., 2000) as well as a particular association between pVAQ and Δ*Glu*_*Sw e*_. These findings may suggest that a strong emotional interference effect was present during the processing of VA-related stimuli. An examination of the changes in Glu in response to nonsocial stimuli, e.g., neutral words, may be worthwhile to explore in a future study to reveal the exact neurochemical response related to VA in the fronto-limbic circuit. Finally, participants in our study were young healthy adults who had low to moderate parental VA experience and clinically subthreshold anxiety or depressive symptoms. As we were able to observe meaningful results from this group of healthy subjects, a further investigation of neurochemical changes using functional spectroscopy on a larger number of subjects with a wide range of VA experience and diverse psychopathology could help understand the neurochemical alterations associated with VA during human brain development.

## Conclusion

We demonstrated neurochemical changes corresponding to the processing of VA-related stimuli in the vmPFC and AMHC using fMRS based on a 3T clinical standard scanner. Individual parental VA severity played a moderating role on Glu responsiveness while processing emotional interference from swear word stimuli. In addition, we found that the change in Glu relative to neuronal density in the vmPFC was a significant predictor for state-trait anxiety as well as depressive mood. Although technical difficulties delimitated the interpretation of the AMHC data and the interrelationship between the two key VOIs investigated, our approach suggests the possibility of using MR spectroscopy to study the underlying mechanism of these emotional processes in the human brain.

## Acknowledgements

We would like to acknowledge Prof. Silvia Mangia at the University of Minnesota on her contribution of discussion throughout the project. This research was supported by the KAIST Venture Research Program for Graduate & Ph.D students, the Brain Research Program through the National Research Foundation of Korea (NRF) funded by the Ministry of Science & ICT (NRF - 2016M3C7A1914448 and NRF-2022M3E5E8081200), Ministry of Health & Welfare of Republic of Korea (KHIDI HI14C1135), and by BK21 Plus Fund (22A20151313464).

